# Signatures of adaptive decreased virulence of deformed wing virus in an isolated population of wild honey bees (*Apis mellifera*)

**DOI:** 10.1101/2022.12.09.519656

**Authors:** Allyson M. Ray, Emma C. Gordon, Thomas D. Seeley, Jason L. Rasgon, Christina M. Grozinger

**Affiliations:** The Pennsylvania State University; Cornell University; Pennsylvania State University; Penn State University

**Keywords:** virulence, deformed wing virus, *Apis mellifera*, *Varroa destructor*, virus evolution, mite-surviving honey bees

## Abstract

Understanding the ecological and evolutionary processes that drive host-pathogen interactions is critical for combating epidemics and conserving species. The *Varroa destructor* mite and deformed wing virus (DWV) are two synergistic threats to Western honey bee (*Apis mellifera*) populations across the globe. Distinct honey bee populations have been found to self-sustain despite *Varroa* infestations, including colonies within the Arnot Forest outside Ithaca, NY, USA. We hypothesized that in these honey bee populations, DWV has been selected to produce an avirulent infection phenotype, allowing for the persistence of both host and disease-causing agents. To investigate this, we assessed the presence and titer of viruses in bees from the Arnot Forest and managed apiaries, and assessed genomic variation and virulence differences between DWV isolates. Across groups, we found viral abundance was similar, but viral genotypes were distinct. We also found that infections with viral isolates from the Arnot Forest resulted in higher survival and lower rates of symptomatic deformed wings, compared to analogous isolates from managed colonies, providing preliminary evidence to support the hypothesis of adaptive decreased viral virulence. Overall, this multi-level investigation of virus genotype and phenotype across different contexts reveals critical insight into global bee health and the ecological and evolutionary processes that drive host-pathogen interactions.

## Introduction

Antagonistic relationships between disease-causing agents, such as pathogens and parasites, and their hosts, are driven by complex interactions modulated by ecological and evolutionary processes (Ebert & Fields, 2020; Retel, Markle, Becks, & Feulner, 2019). Both biotic and abiotic factors can influence disease outcomes and impose selective pressures on both host and pathogen, shaping co-evolutionary dynamics across different contexts (Penczykowski, Laine, & Koskella, 2015). Understanding how these reciprocal exchanges interplay at the genome level is critical for combating epidemics, supporting agricultural systems, and protecting vulnerable species in a changing global climate (Cable et al., 2017; Galvani, 2003).

Population declines in insects broadly, and, in particular, insect pollinator species including the Western honey bee (*Apis mellifera*), have been increasingly documented in recent decades (Hallmann et al., 2017; Potts et al., 2010; Wagner, 2020; Wagner, Grames, Forister, Berenbaum, & Stopak, 2021). Research into honey bee declines has identified multiple factors linked to declining bee health (Goulson, Nicholls, Botías, & Rotheray, 2015). Some factors, as well as their synergistic interactions, include human-driven landscape changes which reduce the flowering plants bees depend on for food, pesticide exposure, climate change, and disease. The dual epidemics of *Varroa destructor* mites and deformed wing virus (DWV) are the primary stressors driving global honey bee colony losses, particularly in temperate regions of the US and Europe (Dainat, Evans, Chen, Gauthier, & Neumann, 2012b). *V. destructor*, an ectoparasite which reproduces on developing bee pupae, expanded it host species from just the Eastern honey bee, *Apis cerana*, to also the Western honey bee, *A. mellifera*, in the last century (Locke, 2016; Traynor et al., 2020). The introduction of *Varroa* to *A. mellifera* not only introduced a novel parasite with no co-evolved resistance, but also introduced a novel transmission route to a historically benign, but now virulent, global pathogen: DWV. Both DWV and *Varroa* have successfully spread to honey bee populations around the world (Wilfert et al., 2016), synergistically undermining honey bee health at multiple levels (Di Prisco et al., 2016; Nazzi et al., 2012).

*Varroa*-mediated DWV transmission leads to increased titers, resulting in enhanced viral disease (Di Prisco et al., 2016; Ray, Davis, Rasgon, & Grozinger, 2021; Ryabov et al., 2014). High levels of DWV lead to deformed wings in adults, reduced activity and ability to contribute to colony tasks, and increased adult mortality (de Miranda & Genersch, 2010; McMahon et al., 2016; Nazzi et al., 2012). This increased mortality leads to reduced colony survival, particularly in the winter months (Dainat, Evans, Chen, Gauthier, & Neumann, 2012a; Dooremalen et al., 2012; Perry, Søvik, Myerscough, & Barron, 2016).

Without management interventions to reduce levels of *Varroa*, most colonies succumb to mite infestations and associated viral infections within 2-3 years (Fries, Imdorf, & Rozenkranz, 2006; Korpela, Aarhus, Fries, & Hansen, 1992; Martin et al., 2012). Indeed, wild, unmanaged honey bee colonies were decimated when *Varroa* was introduced to the US and Europe in the past decades (Jaffe et al., 2010; Kraus & Page, 1995). Recently, though, distinct honey bee populations across the globe have been found to self-sustain and persist despite ubiquitous stressor exposure (Locke, 2016). One such mite-surviving population is located within the Arnot Forest outside Ithaca, NY, USA. These isolated, wild populations located within the Arnot Forest, however, do not demonstrate slowed or reduced mite reproduction (Seeley, 2007) common to other mite-surviving populations (Mondet et al., 2020). Studies have suggested that these wild colonies are smaller in size than managed honey bee colonies, and more likely to swarm (a process of colony reproduction by fission which temporarily ceases brood production): both traits are associated with less brood in the colony and therefore fewer opportunities for mites to reproduce (Seeley, 2017). However, these traits may not be the only factors that are supporting the survival of wild honey bee colonies in the presence of *Varroa* infestation.

How is it possible for these feral bee populations to survive despite the presence of *Varroa* and DWV? While there is evidence for selection on the genome of the Arnot bee populations (Mikheyev, Tin, Arora, & Seeley, 2015), it does not seem to have resulted in significant physiological resistance to mites (Seeley, 2007). It is possible that, rather than selection on the honey bee or the parasitic *Varroa* mite, pathogens including viruses have undergone rapid change to produce an avirulent infection phenotype, allowing for persistence of both host and disease-causing agents. Both mite-resistant populations on the Island of Gotland, Sweden, as well as unmanaged feral bees in PA have been shown to survive high DWV infections levels (Hinshaw, Evans, Rosa, & López-Uribe, 2021; Locke, Forsgren, & De Miranda, 2014). This could be due to virus-tolerant bee genotypes and/or adaptively avirulent virus populations.

It is predicted that in populations where a virus cannot readily infect new hosts, i.e. where population size is small or hosts (i.e. colonies) are far apart, highly virulent pathogens would be selected against, since infected hosts may succumb to the virulent disease prior to transmission to the next host (Brosi, Delaplane, Boots, & De Roode, 2017; Dynes, Berry, Delaplane, Brosi, & De Roode, 2019; Nolan & Delaplane, 2017; Schmid-Hempel, 2011; Seeley & Smith, 2015; Steinhauer & Holland, 1987). Thus, less virulent viruses are expected to have a selective advantage, and persist because their hosts would survive long enough to allow transmission (Steinhauer & Holland, 1987). While lower colony density in managed apiaries is not predicted to dramatically reduce disease prevalence (Bartlett et al., 2019), these conditions may be met in the Arnot Forest, as colonies here are smaller, more spread out, and more apt to swarm than colonies in most managed apiaries (Seeley, 2017). Additionally, if pathogen spread among wild colonies is primarily by vertical transmission (i.e. from parent colony to daughter colony), then this might also select for decreased virulence (Fries & Camazine, 2001; Schmid-Hempel, 2011). Thus, the viral populations circulating within these small, low-density wild populations may have been selected for reduced virulence, allowing them to persist despite lower rates of transmission. Note, however, that increased horizontal transmission (i.e. among unrelated colonies) is predicted to select for increased virulence. Horizontal transmission can occur when bees, *Varroa*, and/or virus-contaminated materials (such as food stores) are moved between colonies by beekeepers, or when bees from different colonies forage together and share viruses on flowers (McMahon, Wilfert, Paxton, & Brown, 2018).

In this study, we investigated whether there is evidence of decreased virulence of viruses found in a population of dispersed, wild colonies compared to populations of crowded, managed colonies. We first assessed the presence and titer of major honey bee viruses in bees sampled from the Arnot Forest, from managed apiaries in adjacent regions in New York and from apiaries in nearby Pennsylvania. These viruses included DWV, the primary virus transmitted by *Varroa*, as well as black queen cell virus (BQCV), a common bee virus not associated with *Varroa* transmission. From a subset of infected bees from Arnot Forest colonies, and from managed colonies, we sequenced DWV genomes and assessed nucleotide differences across these populations to determine if virus isolates were distinct across groups at the nucleotide level. Furthermore, we assessed virulence differences of these DWV isolates in developing honey bees by conducting experimental infections and then measuring pupal and adult mortality as well as other infection phenotypes. Overall, this multi-level analysis of DWV provides initial evidence that selection for decreased DWV virulence may play a role in allowing isolated bee populations to persist despite being parasitized by *Varroa*.

## Methods

### Honey Bee Collections

Bees were collected from 13 sites across three different groups (based on location and management): Arnot Forest (Arnot), New York Managed (NY), and Pennsylvania Managed (PA) (Figure 1). Collections were conducted between September 19 to October 14, 2019, between 10AM and 5PM on sunny, warm (approximately 18-24°C) days. Bees were captured using insect nets, immobilized on dry ice, and then put into 15 mL conical tubes (labeled for site) and kept on dry ice. Bees from managed colonies were collected from hive entrances, preferentially selecting obvious foragers, indicated by pollen-filled corbiculae (n = 2-5 colonies/apiary, 10-15 bees/colony). As it is technically challenging to locate wild colonies and collect at the entrances of their nests, the Arnot Forest bees were collected while they were foraging on flowers (n = 5-12 bees/site). Upon returning from the field, bees were placed at −80°C for long-term storage. Collection details can be found in Supplemental Table 1.

**Figure 1.**
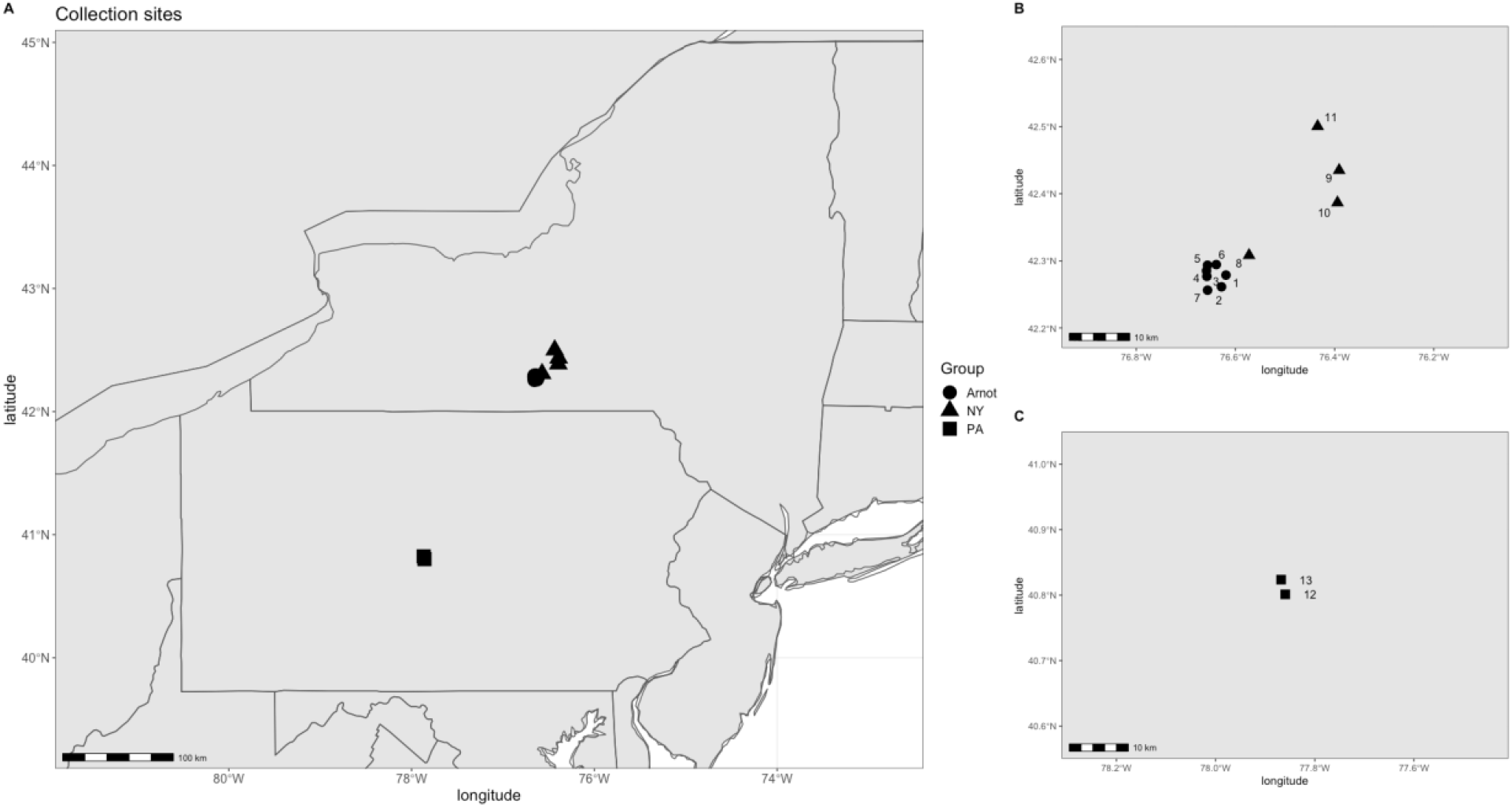
Sampling locations of bees assessed for native DWV infection. (A) The total sites across New York (NY) and Pennsylvania (PA), with a closer view of sites in NY (B) and PA (C). Within each group (distinguished by their point shapes), multiple sites were sampled. For both managed groups (NY and PA), the sites were apiaries from which multiple colonies were sampled.

**Figure 2.**
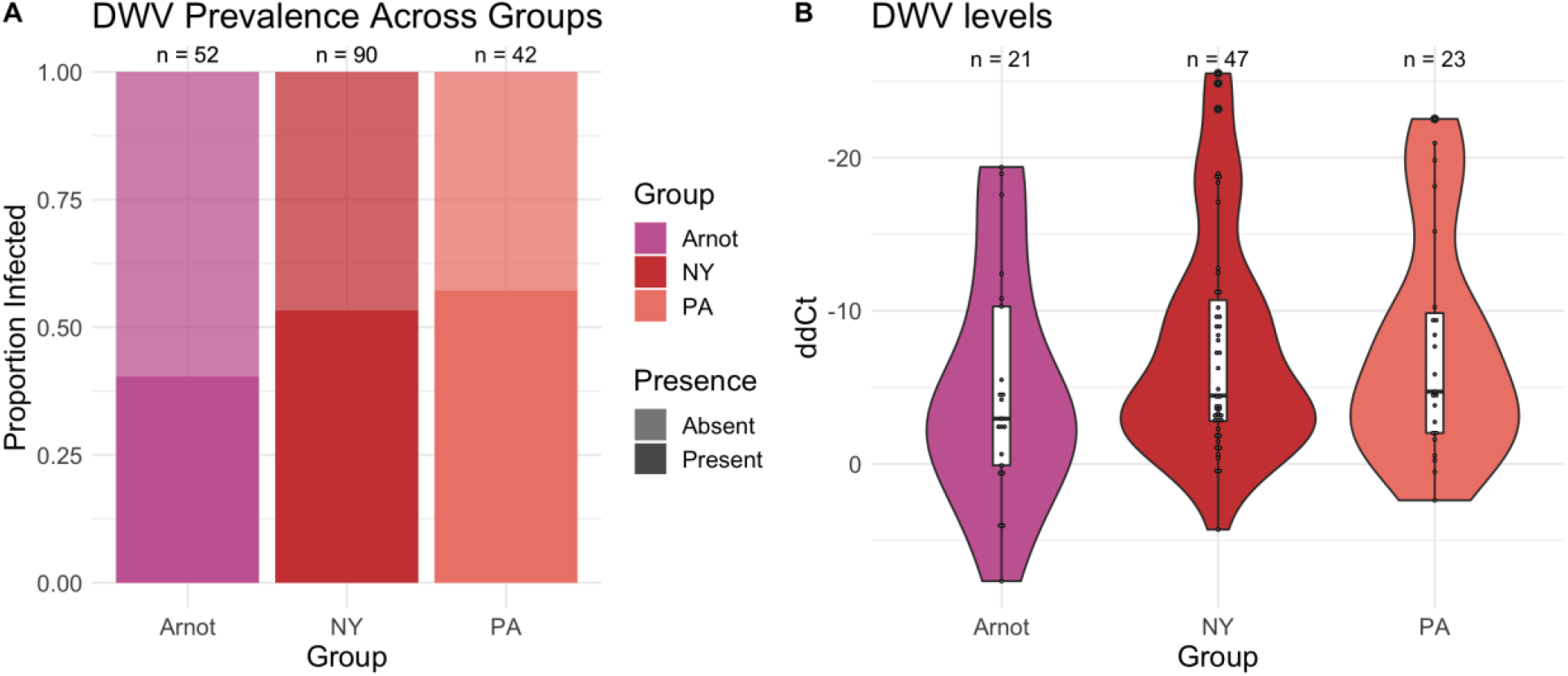
Levels of DWV infection across managed bees and Arnot Forest bees. (A) The proportions of DWV-infected bees did not differ between the bees collected from the Arnot Forest vs. from managed colonies in NY and PA. The rate of infection was around 50% across all groups. Bees were categorized as DWV infected when normalized qPCR dCt was less than 30. (B) Viral loads of DWV-infected bees were similar across groups. Y-axis is reversed, as lower ddCt values are indicative of higher starting template in qPCR reactions.

### Virus Isolation

Viruses were isolated from individual bees as in (Ray et al., 2021). Briefly, bees were homogenized in 500μL of 1xPBS using a Bead Ruptor Elite (Omni International, Kennesaw, GA) at 6.5 m/s for 45 seconds, then centrifuged for 3 minutes at maximum speed (>15,000 x g). Supernatant was passed through a sterile 0.2 μM syringe filter to separate viral particles from honey bee cells, and then was stored at −80°C until RNA purification.

### Virus Quantification by quantitative PCR (qPCR)

RNA was extracted from 30 μL of each virus inoculum using a Direct-zol RNA Miniprep kit (Zymo Research, Irvine, CA) following the manufacturer’s protocol. cDNA was prepared from 200 ng of RNA from each sample using a High-Capacity cDNA Reverse Transcription Kit with RNase Inhibitor (ThermoFisher Scientific, Maltham, MA) following the manufacturer’s protocol. cDNA was diluted 1:20x to allow a sufficient amount of cDNA for all qPCR reactions. qPCR was conducted using PowerUp™ SYBR™ Green Master Mix (ThermoFisher) as in (Ray et al., 2021). Virus was considered “Present” in an individual sample if the normalized mean Ct was < 30. Primers can be found in Supplemental Table 2, and data found in Supplemental Tables 3-9.

### Sequencing and analysis of a subset of isolates

As BQCV was in low abundance across our samples, we focused on DWV for deep sequencing analysis. RNA extracts from a subset of virus isolations with higher DWV levels were submitted to the Pennsylvania State Genomics Core Facility (University Park, PA, USA) for library preparation and sequencing. Libraries were prepared from viral populations of individual bee samples across the Arnot Forest samples as well as the samples from apiaries in New York and Pennsylvania. The 28 samples were sequenced on the Illumina Miseq platform, resulting in 150 nucleotide paired-end stranded mRNA reads. Total reads per sample ranged between 274,942 – 753,035. Reads were assessed for quality with FastQC (version v0.11.9) and quality trimmed with Trimmomatic (version 0.39, option SLIDINGWINDOW:4:25) (Supplemental Table 10).

DWV consensus DWV-A and DWV-B genomes were built using methods described in Ray et al. 2021. Briefly, genomes were created by aligning reads from each sample to DWV-A and -B reference genomes from NCBI (Ref. NC_004830.2 and NC_006494.1, respectively) using Hisat2 (version 2.1.0) (Pertea, Kim, Pertea, Leek, & Salzberg, 2016). Using bcftools (version 1.8) (Li, 2011), variants were called and the consensus fastq sequence files were generated, and from the resulting fasta files, bases with qualities less than 20 were converted to Ns using seqtk (version 1.3-r106) (Li, 2013). DWV levels were low in these samples (Supplemental Table 11), but full-length genomes could be constructed for 10 samples. This resulted in 11 consensus genomes (one sample, 13-1-E was naturally co-infected with DWV-A and DWV-B). Reads were also aligned to a third variant of DWV, variant C, as well as other common bee viruses. However, less than 0.06% of reads within each sample aligned to DWV-C (CEND01000001.1), and less than 0.35% of reads within each sample aligned to other common bee viruses in the USA (acute bee paralysis virus (NC_002548.1), black queen cell virus (NC_003784.1), chronic bee paralysis virus (NC_010711.1), Israeli acute paralysis virus (NC_009025.1), Lake Sinai virus 2 (NC_035467.1), sacbrood virus (NC_002066.1). These viruses were not further examined within the sequence data.

For phylogenetic analyses, multi-sequence alignments of consensus genomes and additional reference genomes (DWV-A reference (NC_004830.2), DWV-B reference (NC_006494.1), and DWV-C (CEND01000001.1)) were generated with Clustal Omega using default settings (version 1.2.3). As the DWV-B genome from Isolate A-3 could not be assembled from the original sequencing, the DWV-B genome constructed from the propagated A-3 isolate from 2021 (see below, “Experimental infection samples and procedure”) was included in its place in the alignment. Multisequence alignment was then imported into MEGAX (version 10.1.8) for Maximum likelihood tree construction with default settings and bootstrapped using 1000 replicates, (Kumar, Stecher, Li, Knyaz, & Tamura, 2018). Consensus genomes, as well as raw sequence reads, will be uploaded to the NCBI Genome and SRA database (accession numbers to be assigned). Variants within DWV-A and -B populations were called using bcftools and annotated using SNPeff (version 5.0) (Cingolani et al., 2012) as in Ray et al. 2021 (Supplemental Tables 12-13).

### Experimental infection samples and procedure

Experimental infections were conducted August – September 2021. Two different colonies (thus representing distinct genotypes) from a Penn State University research apiary were utilized. One colony was headed by a single-drone-inseminated (SDI) queen – this allowed for approximately 75% relatedness between sister bees due to the honey bee’s haplodiploid sex-determination system (Winston, 1991). This minimized the effects of differing honey bee host genetics influencing DWV infection. The other colony was headed by a naturally (i.e. multiply) mated queen. Prior to infection studies, colonies were assessed for viral infection via PCR; there was no or very low indication of common bee viruses. Both colonies were inspected weekly to confirm health status (i.e., no obvious signs of viral disease, and a low parasite load) and to confirm the presence of the original queen.

To reach a sufficiently high titer of viral genotypes to conduct these experiments, inoculums were propagated in pupae collected from a DWV-free (assessed via qPCR) colony (de Miranda et al., 2013). Pupae at the white-eyed stage (14 days post egg laying) were injected with the viral isolates, then collected on dry ice at 4 days post injection (4DPI). Virus was isolated as described above, and aliquoted to minimize the number of freeze-thaws. DWV was quantified (as described above) and assessed for co-infected BQCV and SBV, as well as sequenced to confirm minimal sequence variation from the original, un-propagated isolate (Supplemental Figure 3). Prior to injections, propagated virus isolates were preferentially selected for low co-infection of non-DWV viruses then normalized to two doses: approximately 5×10^6^ genome equivalents per μL and approximately 5×10^2^ genome equivalents per μL. Virus being actively used was kept at 4°C for no longer than 3 days. Re-naming scheme for DWV isolates used in experimental infections can be found in Supplemental Table 14.

Pupal collections and infections were conducted in a UV sterilized hood to minimize contamination by mold and other opportunistic microbes. Virus populations were injected into honey bee pupae at the white-eyed stage. Pupae that showed eye pigmentation (indicating older than 14 days old), melanization (indicating injury during collection), or *Varroa* within their cell were discarded. 1 μL inoculum was injected using a mouth aspirator with an attached 10 μL capillary tube pulled into a needle. Needles were changed between inocula to avoid contamination. To measure colony DWV levels and the effect of the injection itself on DWV levels, control bees (‘Control’, collected from the colony but not manipulated further) and PBS-injected bees (‘PBS-inject’, injected with the saline solution used for the stock viral isolation) were included as controls.

Injected pupae were kept in 48-well plates that were placed in a desiccator at 75% R.H. within an incubator at 34.5°C. Subsets of samples were collected at 3 days post injection (3DPI) to assess viral titers via qPCR. Pupae were monitored daily for mortality, and when nearing the time of eclosion (approximately 7DPI) they were monitored every 8-12 hours for successfully eclosed bees. ‘Successfully’ eclosed bees were identified as ones having normal pigmentation and high mobility, i.e., noticeable movements around their respective wells (Supplemental Table 15-16). Once eclosed, bees were removed from their individual well with sterilized forceps, inspected for deformed wings (Supplemental Table 17), and placed into Plexiglas cages (10 × 10 × 7 cm), split by group (1-7 bees per cage, depending on eclosion rate), noting the time of transfer. Cages were provided 30% sugar water and honey, ad libitum, replenished daily as needed, and placed within an incubator at 34.5°C and approximately 40-60% R.H. Cages with adult bees were monitored for survival daily, and bees that had perished were removed.

### Virus quantifications from experimental infections

RNA was isolated from abdomens from 3 days post injection pupae collected during the experimental infection experiments using the RNeasy Mini Kit (Qiagen, Hilden, Germany) following manufacturers’ protocol including a DNAse incubation step and quantified using a Nanodrop. cDNA synthesis and qPCR were conducted as described above (Supplemental Table 19-25).

### Statistical analyses

Statistical analyses were conducted in R (version 3.6.3) using the ‘stats’ package (R Core Team, 2020). Pearson’s chi-squared test assessed frequency differences in viral presence across groups (Arnot, NY, and PA) and one-way Analysis of Variance (ANOVA) compared viral loads of infected individuals across groups. Differences in viral loads in experimentally infected samples were assessed using two-way ANOVA across Group and Dose. Kaplan-Meier survival analysis was conducted using the ‘survival’ (version 3.4.0) and ‘survminer’ (version 0.4.9) packages.

## Results

### Deformed wing virus presence and loads do not differ between Arnot Forest and managed bee populations

Viruses were isolated from individual bees collected from the Arnot Forest and managed colonies in New York (NY) and Pennsylvania (PA) All groups had detectable DWV and BQCV. Across groups, there was no significant difference in the prevalence of deformed wing virus (DWV), with approximately 40% of samples infected within the Arnot Forest, compared to slightly higher percentages of 53% and 57% in the managed NY and PA samples, respectively (Pearson’s Chi-squared test, p-value = 0.2061, Figure 1a). When comparing viral loads within infected individuals, there was no significant difference in the infection level across groups, with all groups having a range of lowly and highly infected bees (One-way ANOVA, p-value = 0.353, Figure 1b). Both master variants (i.e. strains) *deformed wing virus A* (DWV-A) and *deformed wing virus B* (DWV-B) were found across groups (Supplemental Figure 1).

However, the incidence of BQCV was lower in the bees caught in the Arnot Forest (Pearson’s Chi-squared test, BQCV p-value = 0.001,), as were their viral titers (One-way ANOVA, BQCV p-value < 0.001), relative to the bees collected from managed colonies in NY and PA (Supplemental Figure 2).

### DWV genomes are distinct across groups

Highly infected DWV isolates were subjected to RNA sequencing to identify nucleotide variation across viral genomes. Of the 28 sequenced samples, 10 had sufficient viral titers to allow for reconstruction of full viral genomes, corresponding to 3 DWV-A sequences and 8 DWV-B sequences. Consensus genomes clustered by master variant identity (i.e. DWV-A and DWV-B) in phylogenetic analyses of whole genomes (Figure 3). In the two instances where we had multiple DWV-B isolates collected within the same site (site 7 : A-7-1 and −2, Site 13: PA-13-1 and −2), consensus genomes isolated from the same site also clustered together; otherwise, there is no obvious clustering at the level of geographic location or by group (Figure 3).

**Figure 3.**
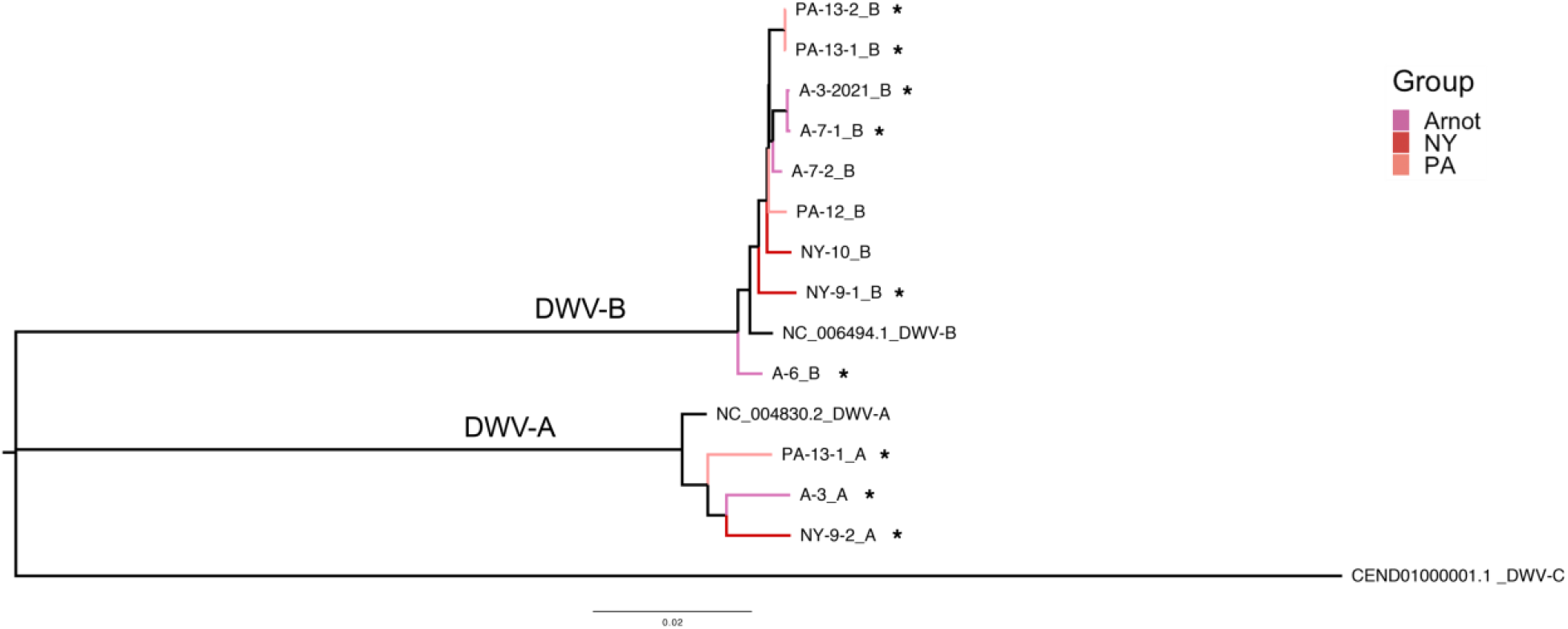
Phylogeny of DWV-A and DWV-B from bees collected in the Arnot Forest (A) and from managed colonies in NY and PA. Maximum likelihood trees with 1000 bootstrap replicates were generated from each isolate’s consensus genome along with reference genomes for DWV-A, DWV-B, and DWV-C (NC_004830.2, NC_006494.1, and CEND01000001.1). Nodes are colored by group. Stars indicate isolates used in experimental infections. As the DWV-B genome from Isolate A-3 was could not be assembled from the original sequencing, the propagated DWV-B from 2021 is shown instead.

Isolates represented primarily by DWV-A show approximately 1.3-1.4% variation across the genome compared to the DWV-A reference genome (NC_004830.2). All isolates had some single nucleotide polymorphism (SNPs) that were unique to each isolate, as well as some that were shared across groups (Figure 4, Supplemental Table 12). DWV-B isolates had about 0.7-0.8% variation compared to the DWV-B reference (NC_006494.1); DWV-B isolates also contained SNPs unique to each isolate, as well as shared within and across groups (Figure 4, Supplemental Table 13).

**Figure 4.**
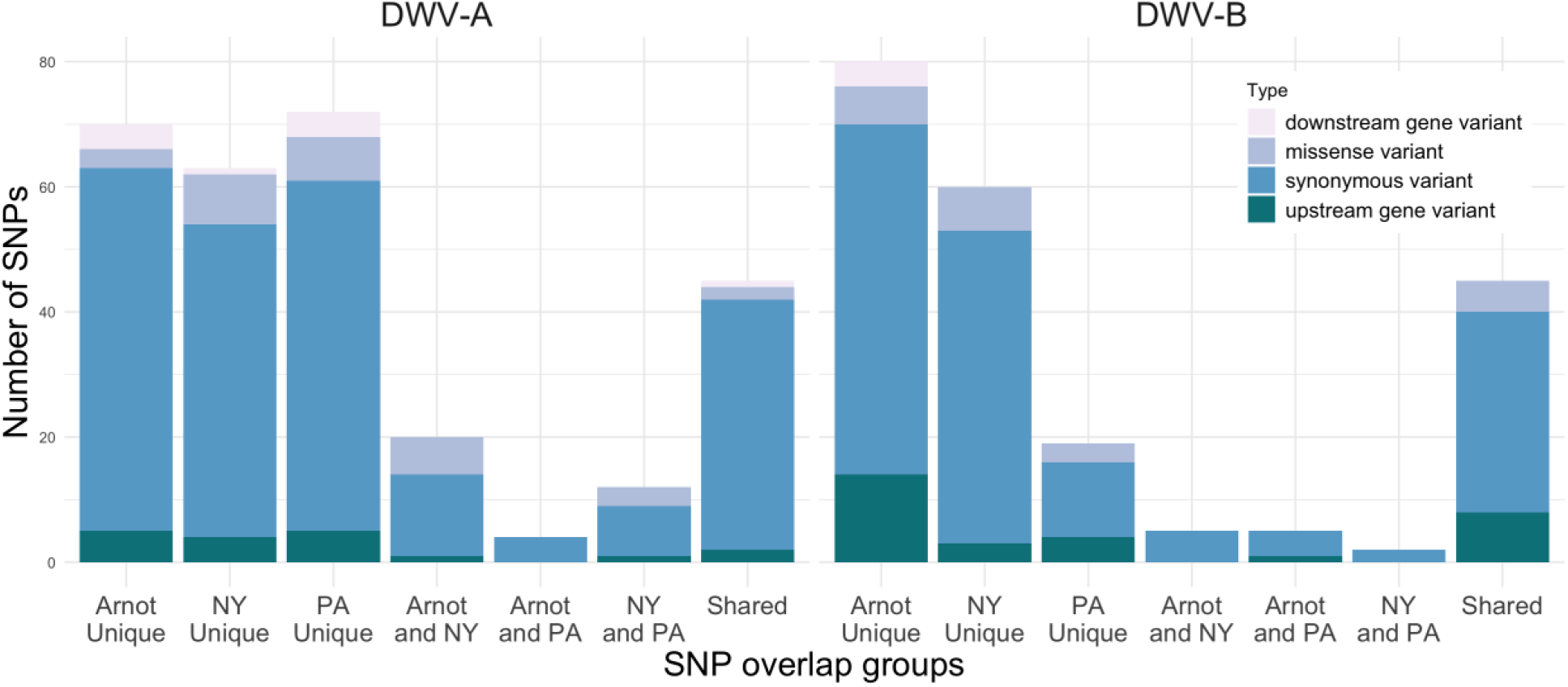
Calculated SNPs across isolates, and determined which were Unique to each group, shared between two groups (e.g. Arnot and NY) or SNPs found across all groups (Shared) for DWV-A (Left) and DWV-B (Right). Type of variant is indicated by color.

Notably, there is one missense variant shared by all 4 Arnot Forest DWV-B isolates: Val896Ile, in the putative capsid protein region of the DWV genome (Supplemental Figure 4). However, overall, SNPs were identified across all groups across the genome, and any missense mutations in DWV-A and DWV-B isolates tended to represent amino acids of the same functional group as the reference allele (Supplemental Tables 12-13).

### Deformed wing virus isolates from Arnot Forest bees were less virulent compared to those from managed bees

Seven DWV isolates, representing a DWV-A and DWV-B from each group, were further assessed for phenotypic differences through experimental infections: 3 isolates from Arnot Forest samples, 2 isolates from NY (from Colony 2 at Site 9), and 2 isolates from PA (from Colony 1 at Site 13). Of these, 3 were found contaminated with other viruses (A-7-1, NY-9-1, NY-9-2 – Supplemental Table 25) and were removed from further analysis. This resulted in a pairwise-comparison of 4 isolates: Arnot vs PA Managed DWV-B (A-6 vs PA-13-2) and Arnot vs PA Managed Mixed (i.e. DWV-A/DWV-B) (A-3 vs PA-13-1).

White-eyed pupae were injected with high doses (approximately 5×10^6^ genome equivalents per μL) or low doses (approximately 5×10^2^ genome equivalents per μL) of an isolate. Others were injected with 1xPBS, as sham-injection controls (PBS). Uninjected pupae were full controls (Control). 3 days post-injection, subsets of pupae were collected to assess infection titers. The remaining bees were allowed to further develop, and later assessed for infection phenotypes including: eclosion rates (i.e. pupal survival rates), rates of symptomatic deformed wings, and adult survival through time.

Viral loads at 3DPI were similar across DWV+ groups, and they were higher than Controls (Supplemental Figure 5). Eclosion rates, i.e. pupal survival, were fairly high (between 71-96%) across all groups and doses (Figure 5a) except for the three groups that had other contaminating viruses, where pupation rates were low (between 0-12%) (Supplemental Table 15); these contaminated groups were subsequently removed from further symptom screening. Interestingly, we also saw more rapid pupation rates across our DWV+ groups compared to controls (Supplemental Figure 6), as has been reported previously (Penn, Simone-Finstrom, Chen, & Healy, 2022).

**Figure 5.**
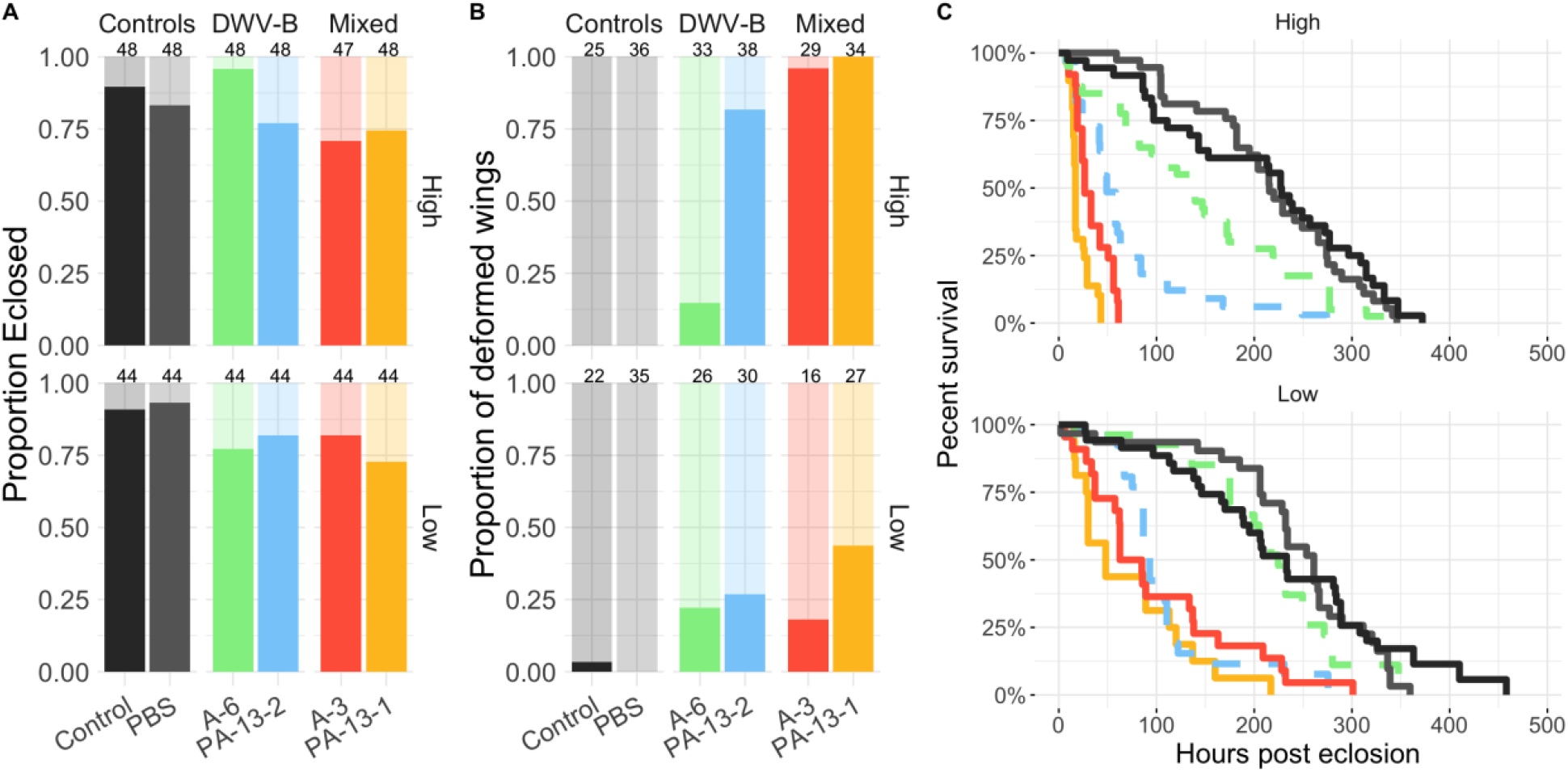
Disease symptoms were less severe with infections from Arnot Forest isolates. Experimental infections were conducted with a pairwise-comparison of 4 isolates: DWV-B Arnot vs PA Managed (A-6 vs PA-13-2) and Mixed Arnot vs PA Managed (A-3 vs PA-13-1). (A) Eclosion percentages were high and similar across groups. (Eclosion status was determined by high mobility around 7DPI.) Percentages of deformed-wing bees were generally lower (B) and the survival percentages were generally higher (C) in bees exposed to Arnot Forest isolates relative to bees exposed to Managed Colony isolates. Samples sizes for eclosion (A) and deformed wing rates (B) can be found above each bar.

Of bees that successfully pupated, those in the DWV+ groups had higher rates of symptomatic deformed wings compared to those that were in the Control group (Figure 5b). Also, mixed groups generally had higher rates compared to DWV-B isolates. When drawing comparisons within the DWV-B isolates, we see that Arnot isolate A-6 had lower rates of deformed wings in the High dose relative to the PA Managed isolate PA-13-2, but that both isolates had similar rates in the Low dose experiments. In the Mixed isolates, again the isolate from the Arnot Forest, A-3, had a lower rate of wing deformities compared to the isolate from the PA Managed group PA-13-1. This difference was most obvious in the Low dose experiments (Figure 5b).

Adult survival over time showed the most distinct disease phenotypes across DWV+ groups (Figure 5c). In the High dose experiments, all the DWV+ groups had significantly lower survival than Controls (PBS and Control), but the Arnot isolate samples consistently had better survival than their managed isolate counterparts. In the Low dose, samples infected with isolate A-6 performed similar to Controls, while the other DWV+ groups again had significantly worse survival than Controls, but were not significantly different from one another (Supplemental Table 26).

## Discussion

We investigated whether adaptive decreased virulence of DWV may contribute to the ability of the feral honey bees of the Arnot Forest to survive. Viral presence and titer were assessed in foragers collected within the Arnot Forest as well as from managed colonies in NY and PA. No significant differences were found between Arnot Forest and managed bees in either their DWV infection rates or their viral loads. However, sequence analyses of DWV isolates revealed unique SNPs associated with the viruses in the different groups of bees. Furthermore, in experimental infections, we found differences—across multiple metrics of virulence—among bees infected with different DWV isolates. Most notably, we found that infections with DWV isolates collected from Arnot Forest bees generally resulted in milder symptoms and better survival compared to infections with DWV isolates collected from managed colonies. Overall, this study provides initial evidence of relatively low virulence of DWV circulating within the Arnot Forest. This is a potential mechanism for colony survival in this forest, despite *Varroa* infestations and pathogen pressure.

By examining individuals, we were able to measure fine-scale infection rates across all groups, and by examining DWV-infected foragers, we began to evaluate which viral genotypes may be circulating in the Arnot Forest. All three groups—workers from wild colonies in the Arnot Forest, and workers from managed colonies in NY or PA—had detectable DWV and BQCV, which shows that the survival of the Arnot Forest bees is not due to a lack of pathogen pressure. We had hypothesized that lower levels of DWV infection might explain the ability of Arnot Forest bees to persist without management, but we found about the same DWV infection rate (approximately 50%) in foragers across all three groups. An ability to suppress DWV titers might also be a honey bee adaptation associated with survival, but we found no evidence of this. The viral loads in infected individuals were similar across all three groups, which is consistent with other studies comparing the viral loads of workers in feral versus managed colonies (Bartlett et al., 2021; Geffre et al., 2021; Hinshaw et al., 2021). Our results suggest that the Arnot Forest bees are instead able to tolerate high levels of infection, as do other bees with mite-resistant genotypes that have demonstrated DWV tolerance (Khongphinitbunjong, de Guzman, et al., 2016; Locke et al., 2014, 2021; Thaduri, Stephan, Miranda, & Locke, 2019).

The infection rates and titers of BQCV were lower in Arnot Forest bees versus managed bees, though the rates we found are still high (78.8% infected, Supplemental Figure 2). BQCV is not associated with vector transmission by *Varroa* (Beaurepaire et al., 2020). BQCV is commonly found in honey bee colonies across the globe (Beaurepaire et al., 2020; Galbraith et al., 2018; Murray et al., 2019), and usually it is not associated with high worker mortality (Chen & Siede, 2007). Therefore, the relatively low infection rates of BQCV may contribute somewhat to Arnot Forest bee survival, but probably it is not the primary basis for bees’ survival. BQCV is readily transferred between bees foraging together in a patch of flowers (Singh et al., 2010), so it is not surprising that it is found in wild colonies. Also, the infection rates of BQCV become high where there are high densities of honey bee colonies (Alger, Burnham, Boncristiani, & Brody, 2019; McNeil et al., 2020). Thus, the lower levels of BQCV in the Arnot Forest bees may be reflect reduced horizontal transmission between foragers from managed and wild colonies, perhaps due to relatively low densities of honey bee colonies.

Both DWV-A and DWV-B genotypes were identified across all three groups. Both master variants, and their recombinants, are virulent (Gisder, Möckel, Eisenhardt, & Genersch, 2018; Natsopoulou et al., 2017; Ryabov et al., 2014), which shows that the survival of the Arnot Forest bees is not due to absence of a particular master variant. At the genome level, consensus sequences of DWV-A and DWV-B did have unique variation across the isolates from the three different groups of colonies, which indicates that there are indeed distinct DWV genotypes circulating in the Arnot Forest. However, consensus DWV-B genomes from the Arnot Forest did not fully cluster with one another, so there does appear to be a “Arnot Forest” sequence variant at the whole-genome level. Similarly, the DWV-A populations in the isolated, mite-resistant colonies in Sweden cluster in the 2009-2010 samples, but not in the 2015 sample (Thaduri, Locke, Granberg, & de Miranda, 2018; Thaduri et al., 2021). Additional sampling over multiple years may reveal more consistent patterns of DWV genotypes within the Arnot Forest.

The individual viral isolates contained a high number of unique SNPs, as well as a high degree of SNPs shared across groups, with the exception of DWV-B isolates from PA. The low amount of variation found in the PA samples is likely due to the fact that both isolates were collected from the same colony, and indeed they show a high number of shared SNPs between one another. A small portion of identified SNPs is shared across all isolates within their group, including a predicted missense variant in the capsid region of the Arnot Forest DWV-B genomes. Mutations in DWV capsid proteins may affect virus cell entry, or recognition by the host (Organtini et al., 2017). Given that this predicted variant results in an amino acid change within the same functional group (valine to isoleucine), it is unclear what affect this SNP may have, if any. Many of the missense variants identified in these populations do not appear to produce a functional change, as no SNPs were identified in putative functional regions and most amino acid substitutions are still within similar functional groups. Nonetheless, sub-consensus and synonymous variation can play important roles in translational efficiency (e.g. codon bias) (Jenkins & Holmes, 2003), RNA secondary structure (Simmonds & Smith, 1999), and pathogen fitness and adaptability (Burch & Chao, 2000), and these may influence the viral dynamics of the Arnot Forest isolates.

While this study presents the first evidence of individual variation in virulence within DWV master variants, due to high pupal mortality in three contaminated groups, we were limited to only assessing four viral isolates through adulthood: two Arnot Forest isolates and two PA managed isolates. Our results provide initial evidence of less virulent DWV populations within bees of the Arnot Forest, or alternatively, more virulent DWV populations in PA managed bees. The isolate with the most distinct infection outcomes, A-6, was also the most diverged DWV-B in the phylogenetic analysis. It is not clear how infection would compare with more genetically similar genotypes, as the other DWV-B isolates we assessed were co-infected, resulting in worsened disease. Moreover, since the A-6 genotype is highly distinct from both its counterpart in the experimental infections (PA-13-2) as well as other Arnot DWV-B genotypes (A-3, A-7-1 and −2), it may therefore not be representative of the sum of Arnot Forest viral population dynamics, per se, and may represent instead a unique variant within the DWV-B classification. To explore adaptive viral avirulence as a mechanism whereby honey bee colonies survive *Varroa* infestations, additional DWV genotypes, from both within and beyond the Arnot Forest, need to be assessed. This could help us to better understand phenotypic variation in infection effectiveness within and across DWV master variants.

The experimental infections reported here provide evidence for DWV adaptation in the Arnot Forest, as well as guidance for future studies in DWV virulence. Overall, the pupation survival rates were comparable for DWV-infected bees and control bees, even at fairly high levels, which has been observed in other studies (Dubois et al., 2020; Tehel et al., 2019). However, the rates of deformed wings and adult survival through time differed among DWV+ groups, indicating the importance of measuring a panel of symptoms during disease phenotyping. We did not find an Arnot Forest isolate that was fully avirulent, although exposure to a Low dose of the Arnot Forest isolate A-6 resulted in adult bee survival that nearly matched that of controls. Samples infected with co-infection isolates (either with both DWV-A and DWV-B, or with DWV and another bee virus) performed worse than samples infected with DWV-B alone. BQCV, which is not naturally transmitted by *Varroa*, has also been shown to be highly virulent when injected directly into the hemolymph of worker bees, simulating *Varroa* transmission (Naggar & Paxton, 2020; Remnant et al., 2019). Furthermore, co-infection of DWV variants can result in increased adult mortality through time, which is consistent with our previous observations of highly virulent DWV-A+DWV-B populations (Ray et al., 2021), and may explain the low rates of DWV co-infection in the naturally infected individuals across all groups (Supplemental Figure 1).

It is important to note that our study tested the impacts of infection with different DWV isolates on honey bees derived from managed stocks. It is possible that the Arnot Forest bees and DWV have co-evolved to be adapted to one another (Lambrechts, Fellous, & Koella, 2006), so there might be even lower virulence in experimental infections of Arnot Forest bees with Arnot DWV isolates. Indeed, honey bee host genotype has been an important factor in DWV infection studies (Khongphinitbunjong, Guzman, et al., 2016; Penn et al., 2022; Ramos-Cuellar et al., 2022), and our study further uncovers how DWV genotype, even within master variant groups, can differ in infection severity. Future studies examining local adaptation (Büchler et al., 2014) and genotype-by-genotype interactions (Barribeau, Sadd, du Plessis, & Schmid-Hempel, 2014; de Roode & Altizer, 2009) will reveal fundamental characteristics of host-pathogen dynamics and avenues for supporting honey bee health.

The relationship among honey bees, *Varroa* mites, viruses, and beekeepers provides a fascinating system in which to study host-pathogen dynamics and evolution (Brosi et al., 2017; McMahon et al., 2018). The introduction of *Varroa* mites provided a novel mechanism for horizontal viral transmission which accelerated the spread of DWV, both within and between colonies, and especially in managed operations (Traynor et al., 2020; Wilfert et al., 2016). There has been considerable focus and interest in selecting for honey bee genotypes that are resistant to both *Varroa* and DWV (Locke, 2016). However, within populations of wild honey bee colonies, decreased opportunities for both horizontal and vertical transmission may result in selection for less virulent viral genotypes (Steinhauer & Holland, 1987), which may be a novel approach to supporting honey bee health. Our study provides the first evidence for this mechanism, and lays the groundwork for further studies examining these dynamics in populations of both managed and wild colonies, and for potentially identifying biomarkers of less virulent DWV populations.

## Acknowledgements

We thank the participants in the Penn State Integrative Pollinator Ecology Colloquium, as well as the members of Grozinger and Rasgon lab for helpful discussions, and Kate Anton for expert beekeeping assistance.

## References

Alger, S. A., Burnham, P. A., Boncristiani, H. F., & Brody, A. K. (2019). RNA virus spillover from managed honeybees (Apis mellifera) to wild bumblebees (Bombus spp.). PLoS ONE, 14(6), 1–13.

Barribeau, S. M., Sadd, B. M., du Plessis, L., & Schmid-Hempel, P. (2014). Gene expression differences underlying genotype-by-genotype specificity in a host–parasite system. Proceedings of the National Academy of Sciences, 111(9), 3496–3501. https://doi.org/10.1073/pnas.1318628111

Bartlett, L. J., Boots, M., Brosi, B. J., Roode, J. C. De, Delaplane, K. S., Hernandez, C. A., & Wilfert, L. (2021). Persistent effects of management history on honeybee colony virus abundances. Journal of Invertebrate Pathology, 179, 1–10. https://doi.org/10.1016/j.jip.2020.107520

Bartlett, L. J., Rozins, C., Brosi, B. J., Delaplane, K. S., Boots, M., Roode, J. C. De, … Boots, M. (2019). Industrial bees: The impact of apicultural intensification on local disease prevalence, (October 2018), 1–11. https://doi.org/10.1111/1365-2664.13461

Beaurepaire, A., Piot, N., Doublet, V., Antunez, K., Campbell, E., Chantawannakul, P., … Dalmon, A. (2020). Diversity and Global Distribution of Viruses of the Western Honey Bee, Apis mellifera. Insects, 11(239), 1–25.

Brosi, B. J., Delaplane, K. S., Boots, M., & De Roode, J. C. (2017). Ecological and evolutionary approaches to managing honeybee disease, 1(September). https://doi.org/10.1038/s41559-017-0246-z

Büchler, R., Costa, C., Hatjina, F., Andonov, S., Meixner, M. D., Conte, Y. Le, … Wilde, J. (2014). The influence of genetic origin and its interaction with environmental effects on the survival of Apis mellifera L. colonies in Europe. Journal of Apicultural Research, 53(2), 205–214. https://doi.org/10.3896/IBRA.1.53.2.03

Burch, C. L., & Chao, L. (2000). Evolvability of an RNA virus is determined by its mutational neighbourhood, 406(August).

Cable, J., Barber, I., Boag, B., Ellison, A. R., Morgan, E. R., Murray, K., … Booth, M. (2017). Global change, parasite transmission and disease control: Lessons from ecology. Philosophical Transactions of the Royal Society B: Biological Sciences, 372(1719). https://doi.org/10.1098/rstb.2016.0088

Chen, Y. P., & Siede, R. (2007). Honey Bee Viruses. Advances in Virus Research, 70(07), 33– 80.

Dainat, B., Evans, J. D., Chen, Y. P., Gauthier, L., & Neumann, P. (2012a). Dead or alive: Deformed wing virus and varroa destructor reduce the life span of winter honeybees. Applied and Environmental Microbiology, 78(4), 981–987. https://doi.org/10.1128/AEM.06537-11

Dainat, B., Evans, J. D., Chen, Y. P., Gauthier, L., & Neumann, P. (2012b). Predictive Markers of Honey Bee Colony Collapse, 7(2). https://doi.org/10.1371/journal.pone.0032151

de Miranda, J. R., Bailey, L., Ball, B. V, Blanchard, P., Budge, G. E., Chejanovsky, N., … van der Steen, J. J. M. (2013). Standard methods for virus research in Apis mellifera. Journal of Apicultural Research, 52(4), 1–56. https://doi.org/10.3896/IBRA.1.52.4.22

de Miranda, J. R., & Genersch, E. (2010). Deformed wing virus. Journal of Invertebrate Pathology, 103(SUPPL. 1), S48–S61. https://doi.org/10.1016/j.jip.2009.06.012

de Roode, J. C., & Altizer, S. (2009). Host – Parasite Genetic Interactions And Virulence-Transmission Relationships In Natural Populations Of Monarch Butterflies. Evolution, 502– 514. https://doi.org/10.1111/j.1558-5646.2009.00845.x

Di Prisco, G., Annoscia, D., Margiotta, M., Ferrara, R., Varricchio, P., Zanni, V., … Pennacchio, F. (2016). A mutualistic symbiosis between a parasitic mite and a pathogenic virus undermines honey bee immunity and health. Proceedings of the National Academy of Sciences, 113(12), 3203–3208. https://doi.org/10.1073/pnas.1523515113

Dooremalen, C. Van, Gerritsen, L., Cornelissen, B., Steen, J. J. M. Van Der, Langeveld, Fr. van, & Blacquière, T. (2012). Winter Survival of Individual Honey Bees and Honey Bee Colonies Depends on Level of Varroa destructor Infestation. PLoS ONE, 7(4). https://doi.org/10.1371/journal.pone.0036285

Dubois, E., Dardouri, M., Schurr, F., Cougoule, N., Sircoulomb, F., & Thiery, R. (2020). Outcomes of honeybee pupae inoculated with deformed wing virus genotypes A and B. Apidologie, 18–34. https://doi.org/10.1007/s13592-019-00701-z

Dynes, T. L., Berry, J. A., Delaplane, K. S., Brosi, B. J., & De Roode, J. C. (2019). Reduced density and visually complex apiaries reduce parasite load and promote honey production and overwintering survival in honey bees. PLoS ONE, 14(5), 1–16. https://doi.org/10.5061/dryad.6j97p92.Funding

Ebert, D., & Fields, P. D. (2020). Host – parasite co-evolution and its genomic signature. Nature Reviews Genetics, 21. https://doi.org/10.1038/s41576-020-0269-1

Fries, I., & Camazine, S. (2001). Implications of horizontal and vertical pathogen transmission for honey bee epidemiology. Apidologie, 32, 199–214.

Fries, I., Imdorf, A., & Rozenkranz, P. (2006). Survival of mite infested (Varroa destructor) honey bee (Apis mellifera) colonies in a Nordic climate. Apidologie, 37, 564–570.

Galbraith, D. A., Fuller, Z. L., Ray, A. M., Brockmann, A., Frazier, M., Gikungu, M. W., … Grozinger, C. M. (2018). Investigating the viral ecology of global bee communities with high-throughput metagenomics. Scientific Reports, (June), 1–14. https://doi.org/10.1038/s41598-018-27164-z

Galvani, A. P. (2003). Epidemiology meets evolutionary ecology. Trends in Ecology & Evolution, 18(3), 132–139.

Geffre, A., Travis, D., Kohn, J., Nieh, J., Geffre, A., Travis, D., … Nieh, J. (2021). Preliminary analysis shows that feral and managed honey bees in Southern California have similar levels of viral pathogens Southern California have similar levels of viral pathogens. Journal of Apicultural Research, 0(0), 1–3. https://doi.org/10.1080/00218839.2021.2001209

Gisder, S., Möckel, N., Eisenhardt, D., & Genersch, E. (2018). In vivo evolution of viral virulence: switching of deformed wing virus between hosts results in virulence changes and sequence shifts, 20, 4612–4628. https://doi.org/10.1111/1462-2920.14481

Goulson, D., Nicholls, E., Botías, C., & Rotheray, E. L. (2015). Bee declines driven by combined Stress from parasites, pesticides, and lack of flowers. Science. https://doi.org/10.1126/science.1255957

Hallmann, C. A., Sorg, M., Jongejans, E., Siepel, H., Hofland, N., Schwan, H., … De Kroon, H. (2017). More than 75 percent decline over 27 years in total flying insect biomass in protected areas. PLoS ONE, 12(10). https://doi.org/10.1371/journal.pone.0185809

Hinshaw, C., Evans, K. C., Rosa, C., & López-uribe, M. M. (2021). The Role of Pathogen Dynamics and Immune Gene Expression in the Survival of Feral Honey Bees. Frontiers in Ecology and Evolution, 8(January), 1–13. https://doi.org/10.3389/fevo.2020.594263

Jaffe, R., Dietemann, V., Allsopp, M. H., Costa, C., Crewe, R. M., DALL’OLIO, R., … Moritz, R. F. A. (2010). Estimating the Density of Honeybee Colonies across Their Natural Range to Fill the Gap in Pollinator Decline Censuses. Conservation Biology, 24(2), 583–593. https://doi.org/10.1111/j.1523-1739.2009.01331.x

Jenkins, G. M., & Holmes, E. C. (2003). The extent of codon usage bias in human RNA v iruses and its e v olutionary origin. Virus Research, 92.

Khongphinitbunjong, K., de Guzman, L. I., Rinderer, T. E., Tarver, M. R., Frake, A. M., Chen, Y., & Chantawan-nakul, P. (2016). Responses of Varroa-resistant Honey Bees (Apis mellifera L.) to Deformed Wing Viru. Journal of Asia-Pacific Entomology, 19(4), 921–927. https://doi.org/10.1016/j.aspen.2016.08.008

Khongphinitbunjong, K., Guzman, L. I. De, Rinderer, T. E., Tarver, M. R., Frake, A. M., Chen, Y., & Chantawannakul, P. (2016). Responses of Varroa-resistant Honey Bees (Apis mellifera L.) to Deformed Wing Virus. Journal of Asia-Pacific Entomology, 19(4), 921– 927. https://doi.org/10.1016/j.aspen.2016.08.008

Korpela, S., Aarhus, A., Fries, I., & Hansen, H. (1992). Varroa jacobsoni Oud. in cold climates: population growth, winter mortality and influence on the survival of honey bee colonies. Journal of Apicultural Research, 31(3–4), 157–164. https://doi.org/10.1080/00218839.1992.11101278

Kraus, B., & Page, R. E. (1995). Effect of Varroa jacobsoni (Mesostigmata: Varroidae) on Feral Apis mellifera (Hymenoptera: Apidae) in California. Environmental Entomology, 24(6), 1473–1480. https://doi.org/10.1093/ee/24.6.1473

Lambrechts, L., Fellous, S., & Koella, J. C. (2006). Coevolutionary interactions between host and parasite genotypes, 22(1). https://doi.org/10.1016/j.pt.2005.11.008

Locke, B. (2016). Natural Varroa mite-surviving Apis mellifera honeybee populations. Apidologie, 47(3), 467–482. https://doi.org/10.1007/s13592-015-0412-8

Locke, B., Forsgren, E., & De Miranda, J. R. (2014). Increased tolerance and resistance to virus infections: A possible factor in the survival of Varroa destructor-resistant honey bees (Apis mellifera). PLoS ONE, 9(6). https://doi.org/10.1371/journal.pone.0099998

Locke, B., Thaduri, S., Stephan, J. G., Low, M., Blacquière, T., Dahle, B., … Miranda, J. R. De. (2021). Adapted tolerance to virus infections in four geographically distinct Varroa destructor-resistant honeybee populations. Scientific Reports, 1–12. https://doi.org/10.1038/s41598-021-91686-2

Martin, S. J., Highfield, A. C., Brettell, L., Villalobos, E. M., Budge, G. E., Powell, M., … Schroeder, D. C. (2012). Global Honey Bee Viral Landscape Altered by a Parasitic Mite, 336(June), 1304–1307.

McMahon, D. P., Natsopoulou, M. E., Doublet, V., Fürst, M., Weging, S., Brown, M. J. F., … Paxton, R. J. (2016). Elevated virulence of an emerging viral genotype as a driver of honeybee loss. Proceedings of the Royal Society B: Biological Sciences, 283(1833). https://doi.org/10.1098/rspb.2016.0811

McMahon, D. P., Wilfert, L., Paxton, R. J., & Brown, M. J. F. (2018). Emerging Viruses in Bees: From Molecules to Ecology. Advances in Virus Research (1st ed.). Elsevier Inc. https://doi.org/10.1016/bs.aivir.2018.02.008

Mcneil, D. J., Mccormick, E., Heimann, A. C., Kammerer, M., Douglas, M. R., Goslee, S. C., … Hines, H. M. (2020). Bumble bees in landscapes with abundant floral resources have lower pathogen loads. Scientific Reports, 10, 1–12. https://doi.org/10.1038/s41598-020-78119-2

Mikheyev, A. S., Tin, M. M. Y., Arora, J., & Seeley, T. D. (2015). Museum samples reveal rapid evolution by wild honey bees exposed to a novel parasite. Nature Communications, 6(7991). https://doi.org/10.1038/ncomms8991

Mondet, F., Beaurepaire, A., Mcafee, A., Locke, B., Alaux, C., Blanchard, S., … Le, Y. (2020). Honey bee survival mechanisms against the parasite Varroa destructor: a systematic review of phenotypic and genomic research efforts. International Journal for Parasitology, 50(6– 7), 433–447. https://doi.org/10.1016/j.ijpara.2020.03.005

Murray, E. A., Burand, J., Trikoz, N., Schnabel, J., Grab, H., & Danforth, B. N. (2019). Viral transmission in honey bees and native bees, supported by a global black queen cell virus phylogeny. Environmental Microbiology, 21(3), 972–983. https://doi.org/10.1111/1462-2920.14501

Naggar, Y. Al, & Paxton, R. J. (2020). Black Queen Cell Virus in Adult Honey Bees, Posing a Future Threat to Bees and Apiculture. Viruses, 12(5), 535. Retrieved from https://doi.org/10.3390/v12050535

Natsopoulou, M. E., McMahon, D. P., Doublet, V., Frey, E., Rosenkranz, P., & Paxton, R. J. (2017). The virulent, emerging genotype B of Deformed wing virus is closely linked to overwinter honeybee worker loss. Scientific Reports, 7(1), 1–9. https://doi.org/10.1038/s41598-017-05596-3

Nazzi, F., Brown, S. P., Annoscia, D., Del Piccolo, F., Di Prisco, G., Varricchio, P., … Pennacchio, F. (2012). Synergistic parasite-pathogen interactions mediated by host immunity can drive the collapse of honeybee colonies. PLoS Pathogens, 8(6). https://doi.org/10.1371/journal.ppat.1002735

Nolan, M. P., & Delaplane, K. S. (2017). Distance between honey bee Apis mellifera colonies regulates populations of Varroa destructor at a landscape scale. Apidologie, 48(1), 8–16. https://doi.org/10.1007/s13592-016-0443-9

Organtini, L. J., Shingler, K. L., Ashley, R. E., Capaldi, E. A., Durrani, K., Dryden, K. A., … Hafenstein, S. L. (2017). Honey Bee Deformed Wing Virus Structures Reveal that Conformational Changes Accompany Genome Release. Journal of Virology, 91(2), 10–13.

Penczykowski, R. M., Laine, A., & Koskella, B. (2015). Understanding the ecology and evolution of host – parasite interactions across scales. Evolutionary Applications, 9(1), 37– 52. https://doi.org/10.1111/eva.12294

Penn, H. J., Simone-Finstrom, M. D., Chen, Y., & Healy, K. B. (2022). Honey Bee Genetic Stock Determines Deformed Wing Virus Symptom Severity but not Viral Load or Dissemination Following Pupal Exposure, 13(June), 1–19. https://doi.org/10.3389/fgene.2022.909392

Perry, C. J., Søvik, E., Myerscough, M. R., & Barron, A. B. (2016). Rapid behavioral maturation accelerates failure of stressed honey bee colonies. Proceedings of the National Academy of Sciences, 113(30). https://doi.org/10.1073/pnas.1610243113

Potts, S. G., Biesmeijer, J. C., Kremen, C., Neumann, P., Schweiger, O., & Kunin, W. E. (2010). Global pollinator declines: Trends, impacts and drivers. Trends in Ecology and Evolution, 25(6), 345–353. https://doi.org/10.1016/j.tree.2010.01.007

R Core Team. (2020). R: A language and environment for statistical computing. R Foundation for Statistical Computing, Vienna, Austria. Retrieved from https://www.r-project.org/

Ramos-Cuellar, A. K., De la Mora, A., Contreras-Escareño, F., Morfin, N., Tapia-Gonzalez, J. M., Macias-Macias, J. O., … Guzman-Novoa, E. (2022). Genotype, but Not Climate, Affects the Resistance of Honey Bees (Apis mellifera) to Viral Infections and to the Mite Varroa destructor. Veterinary Sciences, 9(358). https://doi.org/10.3390/vetsci9070358

Ray, A. M., Davis, S. L., Rasgon, J. L., & Grozinger, C. M. (2021). Simulated vector transmission differentially influences dynamics of two viral variants of deformed wing virus in honey bees (Apis mellifera). Journal of General Virology, 102(11). https://doi.org/10.1099/jgv.0.001687

Remnant, E. J., Mather, N., Gillard, T. L., Yagound, B., Beekman, M., & Beekman, M. (2019). Direct transmission by injection affects competition among RNA viruses in honeybees.

Retel, C., Markle, H., Becks, L., & Feulner, P. G. D. (2019). Ecological and Evolutionary Processes Shaping Viral Genetic Diversity. Viruses, 11(220). https://doi.org/10.3390/v11030220

Ryabov, E. V., Wood, G. R., Fannon, J. M., Moore, J. D., Bull, J. C., Chandler, D., … Evans, D. J. (2014). A Virulent Strain of Deformed Wing Virus (DWV) of Honeybees (Apis mellifera) Prevails after Varroa destructor-Mediated, or In Vitro, Transmission. PLoS Pathogens, 10(6). https://doi.org/10.1371/journal.ppat.1004230

Schmid-Hempel, P. (2011). Evolutionary parasitology: the integrated study of infections, immunology, ecology, and genetics. Oxford University Press.

Seeley, T. D. (2007). Honey bees of the Arnot Forest: a population of feral colonies persisting with Varroa destructor in the northeastern United States, 38, 19–29.

Seeley, T. D. (2017). Life-history traits of wild honey bee colonies living in forests around Ithaca, NY, USA, 743–754. https://doi.org/10.1007/s13592-017-0519-1

Seeley, T. D., & Smith, M. L. (2015). Crowding honeybee colonies in apiaries can increase their vulnerability to the deadly ectoparasite Varroa destructor, 716–727. https://doi.org/10.1007/s13592-015-0361-2

Simmonds, P., & Smith, D. B. (1999). Structural Constraints on RNA Virus Evolution. Journal of Virology, 73(7), 5787–5794.

Singh, R., Levitt, A. L., Rajotte, E. G., Holmes, E. C., Ostiguy, N., VanEngelsdorp, D., … Cox-foster, D. L. (2010). RNA Viruses in Hymenopteran Pollinators: Evidence of Inter-Taxa Virus Transmission via Pollen and Potential Impact on Non-Apis Hymenopteran Species. PLoS ONE, 5(12). https://doi.org/10.1371/journal.pone.0014357

Steinhauer, D. A., & Holland, J. J. (1987). Rapid Evolutionf of RNA viruses. Annual Review of Microbiology, 41(1), 409–431.

Tehel, A., Vu, Q., Bigot, D., Gogol-döring, A., Koch, P., Jenkins, C., … Paxton, R. (2019). The Two Prevalent Genotypes of an Emerging Equally Low Pupal Mortality and Equally High Wing Deformities in Host Honey Bees. Viruses, 11(2) 114, 1–18. https://doi.org/10.3390/v11020114

Thaduri, S., Locke, B., Granberg, F., & de Miranda, J. R. (2018). Temporal changes in the viromes of Swedish varroa-resistant and varroa-susceptible honeybee populations. PLoS ONE, in press, 1–17. https://doi.org/10.1371/journal.pone.0206938

Thaduri, S., Marupakula, S., Terenius, O., Onorati, P., Roth, C. T., Locke, B., & Miranda, J. R. De. (2021). Global similarity, and some key differences, in the metagenomes of Swedish varroa - surviving and varroa - susceptible honeybees. Scientific Reports, (0123456789), 1– 15. https://doi.org/10.1038/s41598-021-02652-x

Thaduri, S., Stephan, J. G., Miranda, J. R. De, & Locke, B. (2019). Disentangling host-parasite-pathogen interactions in a varroa-resistant honeybee population reveals virus tolerance as an independent, naturally adapted survival mechanism, (April), 1–10. https://doi.org/10.1038/s41598-019-42741-6

Traynor, K. S., Mondet, F., Miranda, J. R. De, Techer, M., Kowallik, V., Oddie, M. A. Y., … Mcafee, A. (2020). Varroa destructor: A Complex Parasite, Crippling Honey Bees Worldwide. Trends in Parasitology, 36(7), 592–606. https://doi.org/10.1016/j.pt.2020.04.004

Wagner, D. L. (2020). Insect Declines in the Anthropocene. Annual Review of Entomology, 65.

Wagner, D. L., Grames, E. M., Forister, M. L., Berenbaum, M. R., & Stopak, D. (2021). Insect decline in the Anthropocene: Death by a thousand cuts. Proceedings of the National Academy of Sciences, 118(2), 1–10. https://doi.org/10.1073/pnas.2023989118

Wilfert, L., Long, G., Leggett, H. C., Schmid-Hempel, P., Butlin, R., Martin, S. J. M., … Trueman, J. W. (2016). Deformed wing virus is a recent global epidemic in honeybees driven by Varroa mites. Science (New York, N.Y.), 351(6273), 594–597. https://doi.org/10.1126/science.aac9976

