## Supplemental Figures for "Signatures of adaptive decreased virulence of deformed wing virus in an isolated population of wild honey bees (*Apis mellifera*)"

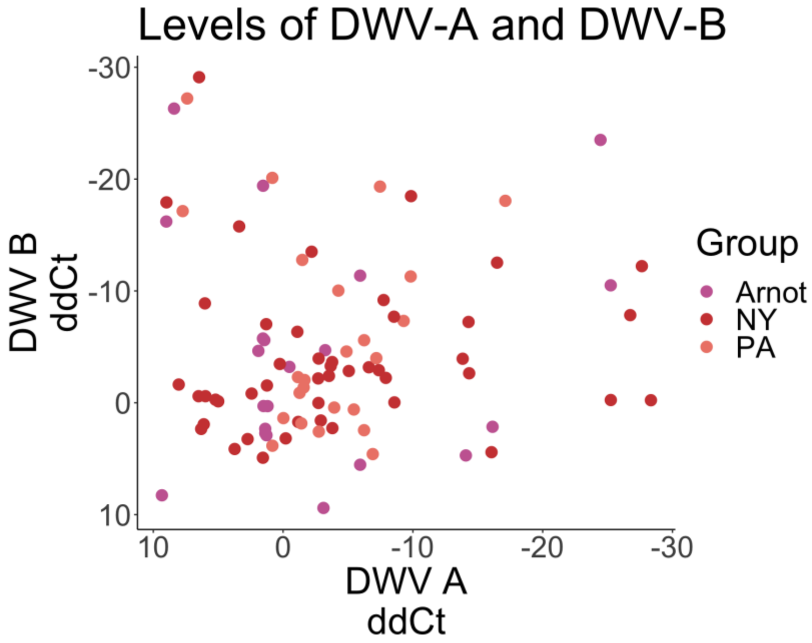


**Supplemental Figure 1 –** levels of DWV-A and DWV-B within DWV-positive samples across groups measured via qPCR with strain specific primers. Each dot represents one individual sample, colored by Group.

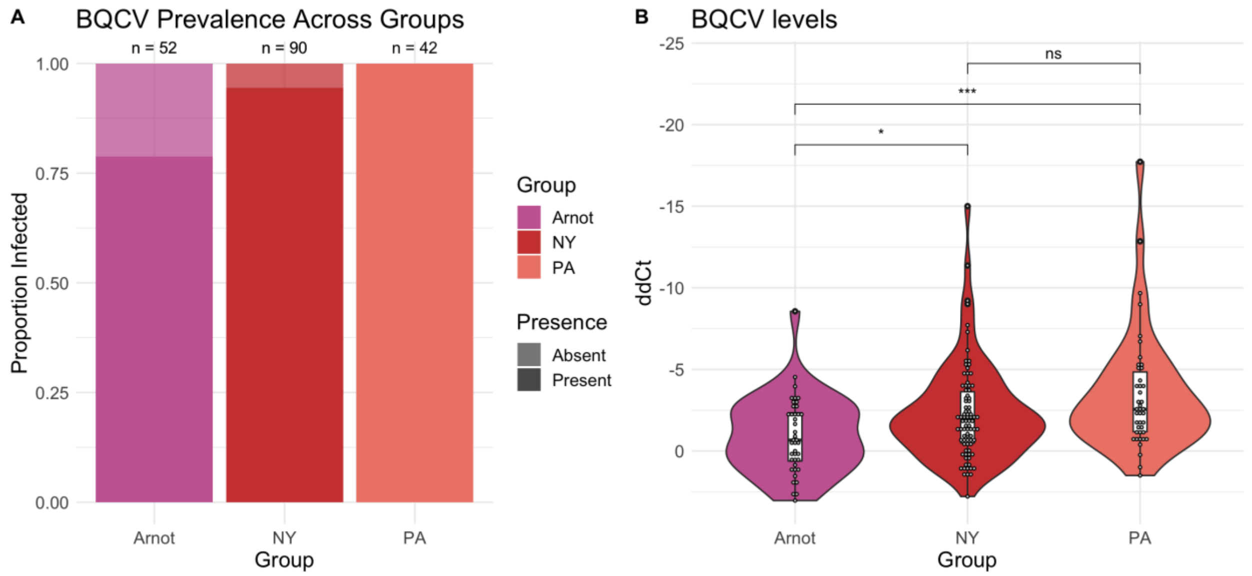


**Supplemental Figure 2 –** (a) Rates of BQCV were significantly lower in bees collected from the Arnot Forest, compared to managed colonies in NY and PA. Infection was determined via a threshold value of less than 30. (b) Of infected samples, BQCV titers within individuals from the Arnot forest were significantly lower than titers from bees collected from managed colonies.

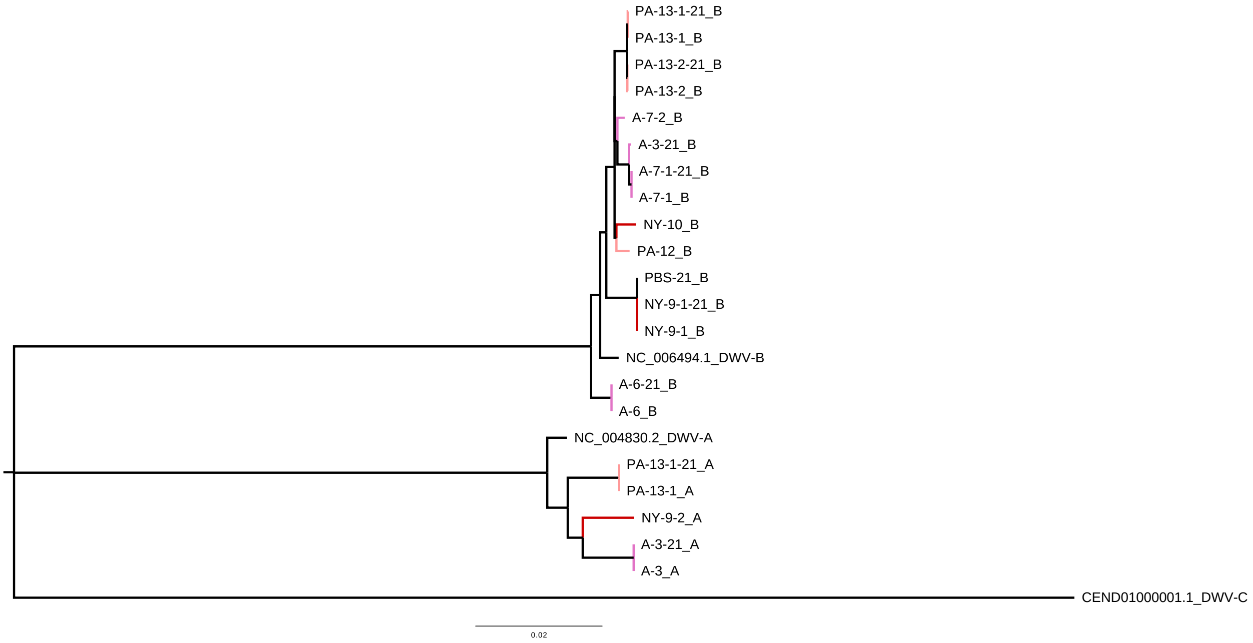


**Supplemental Figure 3 -** Phylogeny of DWV-A and DWV-B from bees collected within the Arnot Forest and managed colonies in NY and PA, in addition to the propagated isolate used in experimental infections (indicated with the suffix "-21"). Maximum likelihood trees with 1000 bootstrap replicates were generated from each isolate's consensus genome along with reference genomes for DWV-A, DWV-B, and DWV-C (NC_004830.2, NC_006494.1, and CEND01000001.1). Nodes are colored by group.

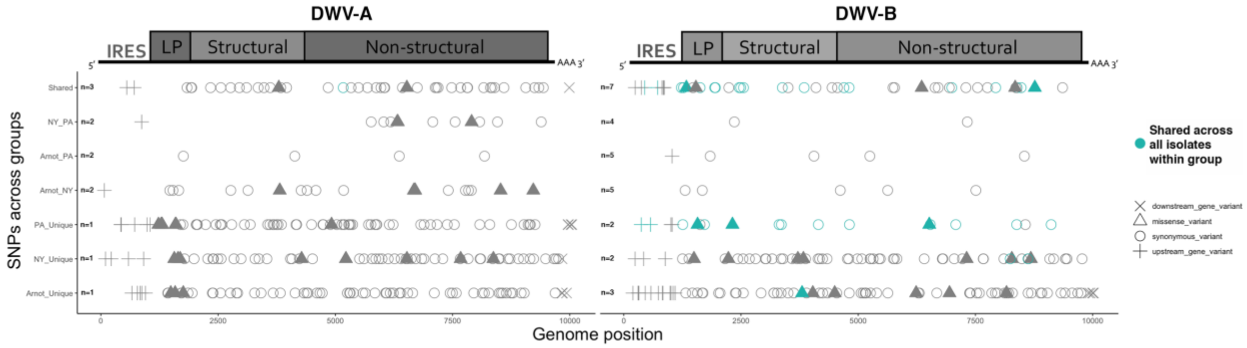


**Supplemental Figure 4** - Location of SNPs for each group across the DWV-A (Left) and DWV-B (Right) genomes. Number of isolates per group is listed near Genome Position 0. Type of variant is indicated by shape. For Groups represented by n ≥ 2 isolates, variants that were shared across ALL individuals within a Group are indicated in Teal.

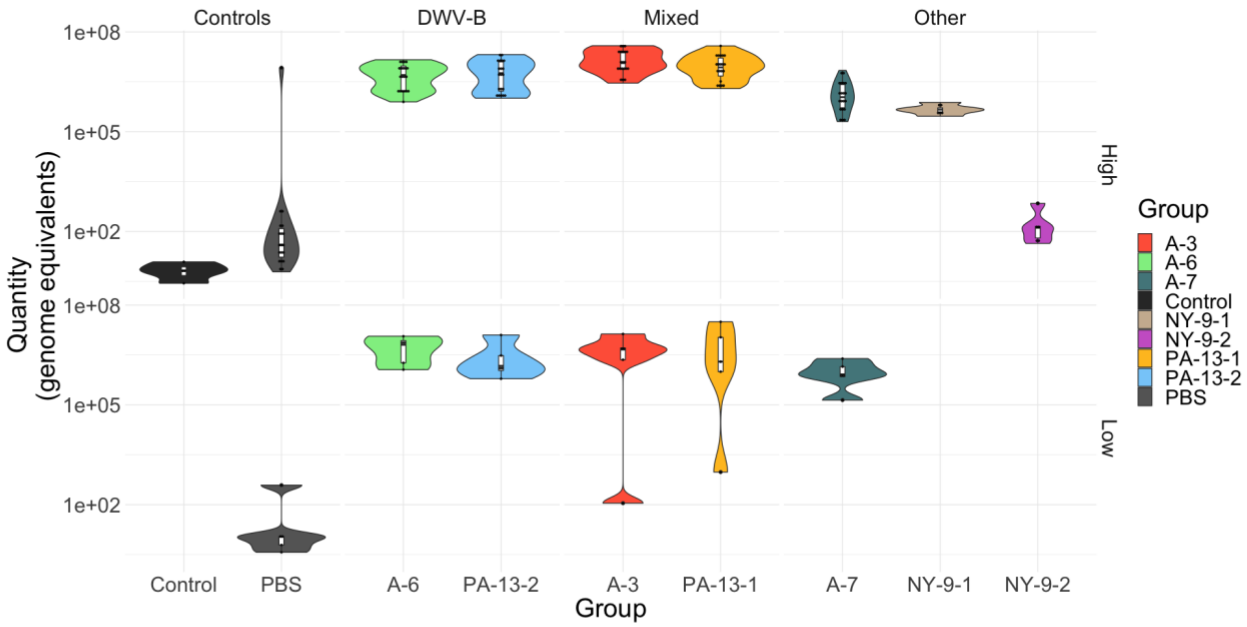


**Supplemental Figure 5** – DWV titers in experimental infections. DWV titers were higher than Controls (Control, PBS) and similar across DWV+ groups within Dose, except for DWV isolate contaminated with other bee viruses (Other : A-7, NY-9-1, and NY-9-2). One exception was one bee in the 'PBS' group at the High dose was naturally infected with DWV, and at similar levels to the experimentally-infected bees. For High dose, n=20 for all groups except the "Others" which had an n=5. Only a subset of groups was assessed for viral loads in the Low Dose – for each group, n= 5, and groups Control, NY-9-1, and NY-9-2 were not assessed.

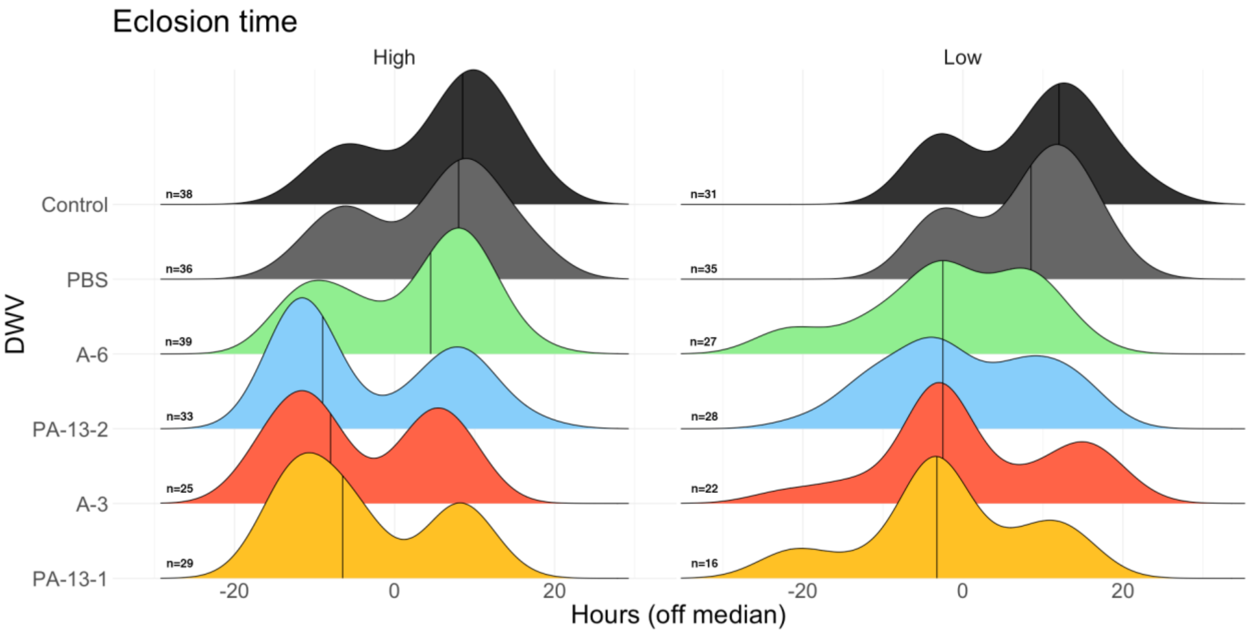


**Supplemental Figure 6** – Distribution of eclosion times of DWV isolates at High and Low doses compared to Controls. DWV+ groups eclose earlier than Control groups across both doses. Sample size (n) per groups is indicated at the left of each distribution, and the line within each plot indicates median value per group. Hours +/- was calculated for each sample within its Trial's median eclosion time (0). Bimodal distributions are likely due to tracking eclosion at 8+ hour intervals, versus tracking eclosion times at shorter intervals (e.g. every 1-2 hours) which may produce smoother distributions.
